# Tbxl represses *Mef2c* gene expression by inducing histone 3 deacetylation of the anterior heart field enhancer

**DOI:** 10.1101/112144

**Authors:** Luna Simona Pane, Filomena Gabriella Fulcoli, Andrea Cirino, Alessandra Altomonte, Rosa Ferrentino, Marchesa Bilio, Antonio Baldini

## Abstract

The *TBX1* gene is haploinsufficient in the 22q11.2 deletion syndrome (22q11.2DS), and genetic evidence from human patients and mouse models points to a major role of this gene in the pathogenesis of this syndrome. Tbx1 can activate and repress transcription and previous work has shown that one of its functions is to negatively modulate cardiomyocyte differentiation. Tbx1 occupies the anterior heart field (AHF) enhancer of the *Mef2c* gene, which encodes a key cardiac differentiation transcription factor. Here we show that increased dosage of *Tbx1* is associated with down regulation of *Mef2c* expression and reduced acetylation of its AHF enhancer in cultured mouse myoblasts. Consistently, 22q11.2DS-derived and in vitro differentiated induced pluripotent stem cells (hiPSCs) expressed higher levels of *Mef2c* and showed increased AHF acetylation, compared to hiPSCs from a healthy donor. Most importantly, we show that in mouse embryos, loss of *Tbx1* enhances the expression of the Mef2c-AHF-Cre transgene in a specific region of the splanchnic mesoderm, and in a dosage-dependent manner, providing an *in vivo* correlate of our cell culture data. These results indicate that Tbx1 regulates the Mef2c-AHF enhancer by inducing histone deacetylation.

## Introduction

During development, cardiac progenitors of the second heart field (SHF) are recruited or incorporated into the cardiac outflow tract and right ventricle (Kelly, Brown and Buckingham, 2001; Mjaatvedt *et al*., 2001; Waldo *et al*., 2001). During this process, cells activate a differentiation program that leads to expression of specialized proteins such as contractile proteins necessary for cardiomyocyte function. This process, still to be dissected in detail, should be tightly regulated so that a sufficient number of progenitors are allowed to proliferate and are prevented from differentiating prematurely. These two basic functions (pro-proliferative and anti-differentiative) are likely effected by a combination of signals and transcription factors, among which Tbx1 is a major candidate (Jerome and Papaioannou, 2001; Lindsay *et al*., 2001; Merscher *et al*., 2001; Rana *et al*., 2014). Indeed, Tbx1 is expressed in cardiac progenitors of the SHF, is shut down as progenitors differentiate, and it regulates positively pro-proliferative signals, and, negatively, pro-differentiative factors such as *Gata4* (Liao *et al*., 2008; Pane *et al*., 2012), *Mef2c* (Pane *et al*., 2012), SRF (Chen *et al*., 2009), and Smad1 (Fulcoli *et al*., 2009). Down regulation of Srf and Smad1 signaling are effected through non-transcriptional mechanisms, but *Mef2c* gene expression appears to be transcriptionally repressed by Tbx1.

Tbx1 regulates its target genes by interacting with the SWI-SNF-like BAF complex and with histone methyltransferases (Stoller *et al*., 2010; Chen *et al*., 2012; Fulcoli *et al*., 2016), reviewed in (Baldini, Fulcoli and Illingworth, 2017). However, there are a number of questions to be answered. In this work, we addressed the question as to how Tbx1 represses *Mef2c* expression. We used three different model systems: a) a mouse myoblast cell line, b) transgenic mouse embryos, and c) differentiating human induced pluripotent stem cells (hiPSCs) from a 22q11.2DS patient and a healthy donor. Results indicate that transcriptional repression is associated with reduced histone 3 acetylation in the region bound by Tbx1 in cultured cells, that is the previously defined AHF enhancer of the *Mef2c* gene (Dodou *et al*., 2004). In addition, we show that *in vivo*, Tbx1 regulation of the AHF enhancer is dosage-dependent and is regionally restricted, suggesting that there are crucial interactions with other transcription factors in regulating this enhancer.

## Materials and Methods

### Generation of human induced pluripotent stem cells (hiPSCs)

Neonatal skin fibroblasts from a 22q11.2DS/DiGeorge syndrome patient were obtained from the Coriell Institute (GM07215); control skin fibroblasts were obtained from an unrelated age-and gender-matched anonymous, healthy donor.

For iPSC induction, retroviruses encoding the human *OCT3/4, SOX2, KLF4, and c-MYC* factors were independently produced by transfecting HEK293T cells with pMXs vectors (Addgene plasmids 17217, 17218, 17219, and 17220) and the combination of moloney gag-pol plasmid pUMVC (Addgene plasmid 8449) and VSV envelope plasmid (Addgene plasmid 8454) in DMEM containing 10% FBS using Fugene HD (Roche). Viral supernatants were harvested after 48 and 72 h, filtered through a 0.45-μm low-protein-binding cellulose acetate filter, concentrated by a spin column (Millipore, Billerica, MA, USA), and used directly to infect twice (24 h apart) 1.5×10^5^ human primary skin fibroblasts (PSFs) in the presence of 8 μg/ml polybrene. After 6 days, cells were seeded on mouse embryonic fibroblast (MEF) feeders at the density of 5×10^4^ cells/10-cm dish and cultured for 4 additional weeks in human ES cell medium, consisting of DMEM/F12 supplemented with 20% knockout serum replacement (KSR, Invitrogen), 2mM L-glutamine, 0.1mM nonessential amino acids, 0.1mM β-mercaptoethanol, 50U/ml penicillin, 50mg/ml streptomycin and 10ng/ml human b-FGF (R&D), before hiPSC colonies were manually picked. Karyotyping of the iPSC lines was performed at the Cell Culture Facility of the Telethon Institute of Genetics and Medicine in Naples, Italy, using standard methods.

### Cardiac differentiation of hiPSCs

To generate hiPSC embryoid bodies (EBs), hiPSC colonies were dissociated into clumps using PBS containing 2.5 mg/ml trypsin (USB, Staufen, Germany), 1 mg/ml collagenase IV (Invitrogen), 20% KSR, and 1 mM CaCl_2_ (10 min at 37°C) and maintained for 3 days in MEF-conditioned human ES medium in low attachment plates. For spontaneous differentiation, medium was then replaced with DMEM/F12 supplemented with 20% FBS, 2 mM L-glutamine, 0.1 mM nonessential amino acids, 0.1 mM β-mercaptoethanol, 50 U/ml penicillin, and 50 *μ*g/ml streptomycin, and EBs were analyzed at day 15 for expression of marker genes of the 3 different germ layers. To improve cardiac differentiation, ascorbic acid (50 μg/ml) was added to the medium, and EBs were plated at day 7 on gelatin-coated dishes for better detection of beating foci.

For induction of human cardiac progenitors, hiPSCs were seeded on MEF feeders and, 1 day later, treated for 4 days with 10 ng/ml human BMP2 (R&D) and 1 μM SU5402 FGF receptor inhibitor (Calbiochem, Darmstadt, Germany) in RPMI 1640 medium supplemented with 2 mM L-glutamine and 2% B27 supplement without vitamin A (Invitrogen), as described previously (Leschik *et al*., 2008; Moretti *et al*., 2010). Conversion of human cardiac progenitors into myocytes and vascular cells (endothelial and smooth muscle cells) was induced by supplementing the culture medium with 50 μg/ml ascorbic acid and 10 ng/ml human VEGF (R&D System).

### Immunofluorescence and alkaline phosphatase activity assay

hiPSCs (undifferentiated or differentiated) were fixed in 3.7% (vol/vol) formaldehyde and subjected to immunostaining by using the following primary antibodies: human Nanog (rabbit polyclonal, Abcam, 1:500), TRA1-81-Alexa-Fluor-488-conjugated (mouse monoclonal, BD Pharmingen, 1:20). Alexa-Fluor-488, -594, and -647 conjugated secondary antibodies specific to the appropriate species were used (Life Technologies, 1:500). Nuclei were detected with 1 μg/ml Hoechst 33528. Direct alkaline phosphatase activity was analyzed using the NBT/BCIP alkaline phosphatase blue substrate (Roche), according to the manufacturer’s guidelines. Microscopy was performed using the imaging systems (DMI6000-AF6000), filter cubes and software from Leica microsystems. Images were pseudo-colored using Adobe Photoshop.

### Mouse lines and immunofluorescence

Mef2c-AHF-Cre mice (Verzi *et al*., 2005) were crossed with *Tbx1*^*/lacZ^ mice (Lindsay *et al*., 2001) to obtain Mef2c-AHF-Cre;Tbx1^+/lacZ^ animals, which were then crossed with *Tbx1*^*flox*/flox^ (Xu *et al*., 2004) or *Tbx1*^*/lacZ^ mice. Tbx1^+/lacZ^ (also referred to as *Tbxl^·/-^*) mice where also crossed with WT mice to harvest embryos to be used for *Tbx1* gene expression by β-galactosidase assay. Animal studies were carried under the auspices of the animal protocol 257/2015-PR (licensed to the AB lab) reviewed, according to Italian regulations, by the Italian Istituto Superiore di Sanità and approved by the Italian Ministero della Salute. Pregnant females (mostly 3-6 months-old) at plug day (E) 9.5 were sacrificed using CO_2_ inhalation. The laboratory applies the ″3Rs″ principles to minimize the use of animals and to limit or eliminate suffering.

Immunofluorescence on paraffin-embedded E9.5 embryo sections was performed using an anti Cre antibody (Novagen, 69050-3, 1:1000), and an anti Isl1 antibody (Hybridoma Bank, 39.4D5-s, 1:50). β-galactosidase assays were performed on whole-mount embryos using salmon-gal (6-Chloro-3-indolyl-beta-D-galactopyranoside, Alfa Aesar, B21131).

### Quantitative reverse transcription PCR (qRT-PCR)

Total mRNA was isolated from PSF, hiPSCs, EBs, and cardiac cells using the Stratagene Absolutely RNA kit and 1*μ*g was used to synthesize cDNA using the High-Capacity cDNA Reverse Transcription kit (Applied Biosystems). Gene expression was quantified by qRT-PCR using 1*μ*l of the RT reaction and the Power SYBR Green PCR Master Mix (Applied Biosystems). Gene expression levels were normalized to GAPDH. A list of primers is provided in Supplementary Tab. 1.

### Cell lines, plasmids and transfections

Mouse C2C12 myoblasts (ATCC, # CRL-1772) were cultured in Dulbecco’s modified Eagle’s medium supplemented with 10% fetal bovine serum. Cultures were tested for mycoplasma infection and were negative. Authentication has been performed using differentiation marker analyses. For differentiation and transient transfection, cells were plated at a density of 1.5×10^5^ cells/well on a 35-mm tissue culture dish and incubated at 37°C in 5% CO_2_. 24 h later, the medium was replaced with a differentiation medium containing 2% horse serum (Hyclone). Transfection of 1 *μ*g of Tbx1-3HA (Chen *et al*., 2012) was performed with X-tremeGENE (Roche, Penzberg, Germany) according to the manufacturer’s instructions.

### Chromatin immunoprecipitation

C2C12 cells and hiPSCs were cross-linked with 1% formaldehyde for 15 min at room temperature and glycine was added to stop the reaction to a final concentration of 0.125 M for 5 min. The cell pellet was suspended in 6 x volumes of cell lysis buffer (10 mM HEPES, 60 mM KCl, 1 mM EDTA, 0.075% v/v NP40, 1 mM DTT and 1X protease inhibitors, adjusted to pH 7.6) in a 1.5 mL tube incubating on ice for 15 min. Isolated nuclei were suspended in Buffer B (LowCell ChIP Kit reagent) and chromatin was sheared into 200–500 bp long fragments using the Covaris S2 Sample Preparation System (Duty Cycle: 5%, Cycles: 5, Intensity: 3, Temperature: 4°C, Cycles per Burst: 200, Power mode: Frequency Sweeping, Cycle Time: 60 seconds, Degassing mode: Continuous). Soluble chromatin was incubated with 6*μ*g of Anti-acetyl-Histone H3 (Millipore, #06-599), anti H3K27Ac (Abcam, ab4729), or normal rabbit/mouse IgG (Santa Cruz Biotechnology, #2027). Following steps included extensive washes and reverse crosslinking following the LowCell ChIP Kit instructions. For quantitative ChIP, we performed real-time PCR of the immunoprecipitated DNA and inputs, using the FastStart Universal SYBR Green Master kit (Roche) and the 7900HT Fast Real-Time PCR System or Step-one plus (Applied Biosystems) using primers specific for the AHF-enhancer region of *Mef2c* (Supplementary Tab. 1).

For ChIP western experiments, chromatin from C2C12 cells (transfected with a 3xHA-tagged mouse Tbx1 transfection vector) was immunoprecipitated using anti HA antibodies (Roche, clone 12CA5, catalog 11583816001) and washed as indicated above. 4x reducing SDS Loading Dye (500 mM Tris HCl, pH 6.8, 8% SDS, 40% glycerin, 20% β-mercaptoethanol, 5 mg/ml bromophenol blue) was added to the beads as well as to the lysates and the supernatants and samples were incubated at 99°C for 20 min and loaded onto SDS-polyacrylamide gel electrophoresis, proteins were then transferred into PVDF membranes. Western blotting was performed using anti-HA (control) and anti HDAC1 antibodies (Abcam, Ab7028).

*Tbx1* knock down in C2C12 cells was obtained by transfection with non-targeted or *Tbx1*-targeted siRNA (Life Technologies, pool of s74767 and s74769, 50nmol/L) using Lipofectamine RNA iMAX (Life Technologies). Cells were cross-linked and processed for ChIP 48 hrs after transfection, as described above.

### Co-immunoprecipitation

C2C12 cells were transfected with aTbx1:3xHA expression vector (1*μ*g) and harvested after 24 hrs for protein extraction. 100 *μ*l of protein-G Dynabeads (Life Technology, #10004D) were cross-linked to 20 *μ*g of anti-HA antibodies (Roche, clone 12CA5, catalog 11583816001) or Control IgG (Abcam, #ab46540). Nuclear extracts were incubated overnight at 4°C with crosslinked antibodies and extensively washed with phosphate-buffered saline (PBS)-0.02% Tween-20. Two consecutive elutions were performed with 0.1M Glycine pH2.8 and immediately neutralized with Tris-HCl pH 8.0. Samples were subjected to SDS polyacrylamide gel electrophoresis and proteins were transferred into PVDF membrane for western blotting analyses with anti-HDAC1 antibody (company, catalog#, dilution), anti-HA antibody (Abcam, Ab7028, 1:200), and VeriBlot for IP secondary antibody (HRP) (Abcam, #ab131366, 1:200).

### Statistical analysis

All data were expressed as means ± s.e.m. (or SD, as indicated in the Figure legends) from independent experiments. Differences between groups were examined for statistical analysis using a 2-tailed Student’s *t* test. Values of *P* < 0.05 were considered significant.

## Results and Discussion

### Increased Tbx1 expression is associated with reduced histone 3 acetylation of Mef2c AHF enhancer sequence

To understand the mechanisms by which Tbx1 represses *Mef2c*, we used mouse myoblast C2C12 cells undergoing muscle differentiation. We transfected a Tbx1 expression vector in these cells at day 0 and let them differentiate (Fig. 1a). The *Mef2c* gene was strongly up-regulated at day 3 of differentiation in control cells, but in *Tbx1*-transfected cells this up-regulation was strongly reduced (Fig. 1b). To exclude that this effect may be due to a delayed differentiation caused by early *Tbx1* transfection, we repeated the experiment by transfecting Tbx1 after the initiation of differentiation, at day 2, and assayed *Mef2c* expression at day 3. Also in this case, we observed a strong reduction of *Mef2c* expression (Fig. 1c).

**Figure 1.**
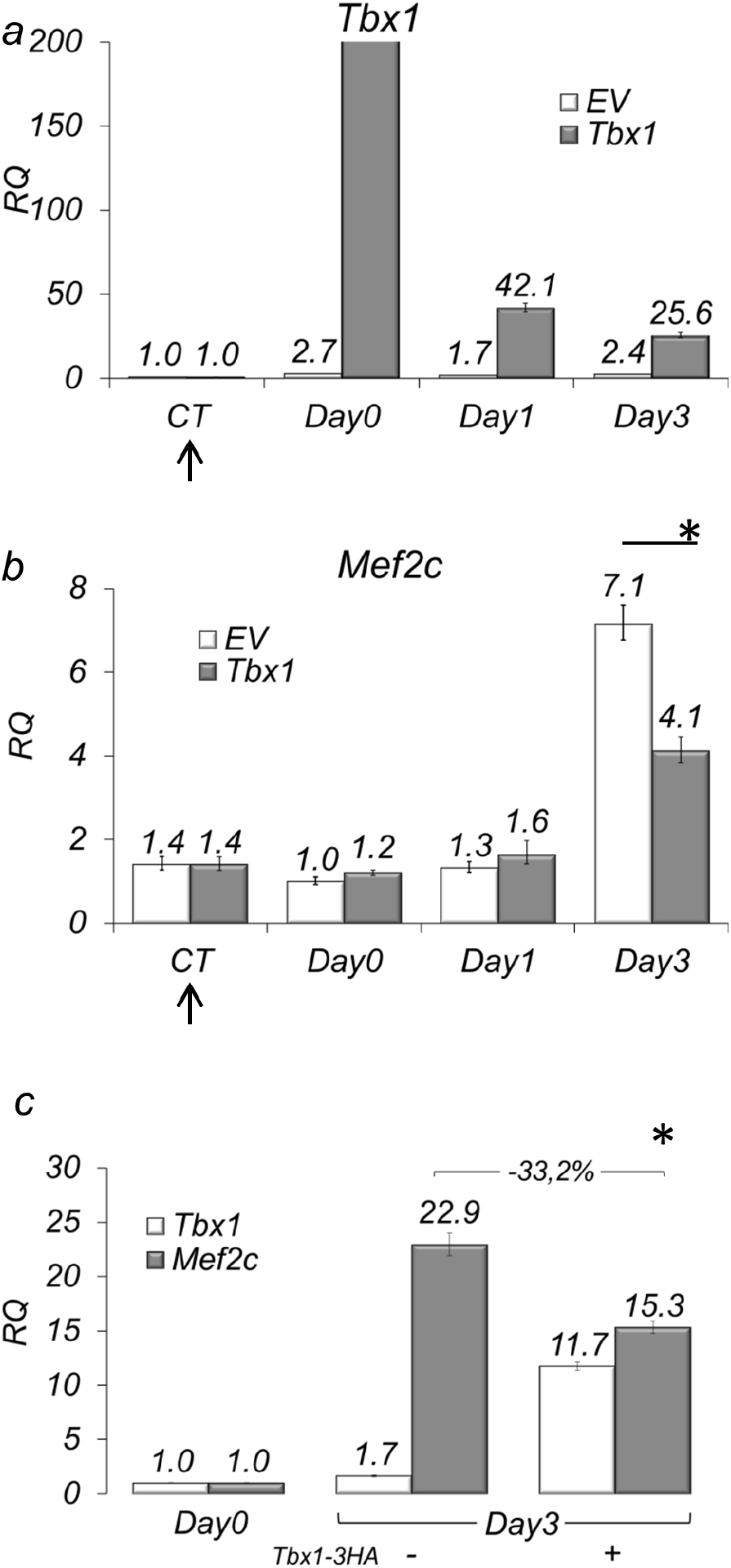
Tbx1 negatively regulates Mef2c expression in C2C12 at Day3. Time course of *Tbx1* (a) and *Mef2c* (b) expression during in vitro C2C12 myoblast differentiation with or without transient transfection of *Tbx1*. Note that Tbx1 negatively regulates *Mef2c* expression at day 3 of differentiation (b). (c) *Mef2c* mRNA levels are affected by *Tbx1* overexpression 24 hours after transfection (performed at day 2). Values are from 3 experiments (mean± S.D.). Asterisks indicate statistically significant difference (P-value less than 0.05). The arrows in panels a-c indicate the time of transfection (24 hours before induction of differentiation, which is on day 0).

Tbx1 ChIP-seq data obtained on P19Cl6 cells (Fulcoli *et al*., 2016) and previously published q-ChIP data on C2C12 cells (Pane *et al*., 2012) revealed enrichment at the *Mef2c* anterior heart field (AHF) enhancer, as previously defined (Dodou *et al*., 2004) (Fig. 2a). In particular, by comparing Tbx1 ChIP-seq data on P19Cl6 cells with histone modification data during cardiac differentiation of mouse embryonic stem cells (ESCs) (Wamstad *et al*., 2012), we noted that the Tbx1-enriched region is relatively poor in H3K27Ac in ESCs but it progressively becomes H3K27Ac-rich during cardiac differentiation. Using available data from ENCODE, we also noted that the same region is acetylated in E14.5 in heart, where *Tbx1* is not expressed (Fig. 2a). These observations suggest that in this region H3 acetylation increases as differentiation progresses. Thus, we reasoned that Tbx1, directly or indirectly, might negatively regulate acetylation in this region. To test this possibility, we performed quantitative chromatin immunoprecipitation (q-ChIP) with an antibody against H3 acetylation (H3-Ac) in C2C12 cells. Results showed that *Tbx1* overexpression substantially reduced H3-Ac enrichment at the *Mef2c* AHF enhancer (Fig. 2b). Consistent results were obtained using antibodies against H3K27Ac on C2C12 cells after *Tbx1* knock down, which increased H3K27Ac enrichment (Supplementary Fig. 1). We addressed the question as to whether the H3 deacetylating effect of Tbx1 may be due to a direct interaction with HDAC1. However, repeated co-IP experiments failed to demonstrate HDAC1-Tbx1 co-immunoprecipitation, suggesting that deacetylation is an indirect effect or that Tbx1 interacts with other HDACs (Supplementary Fig. 2).

**Figure 2.**
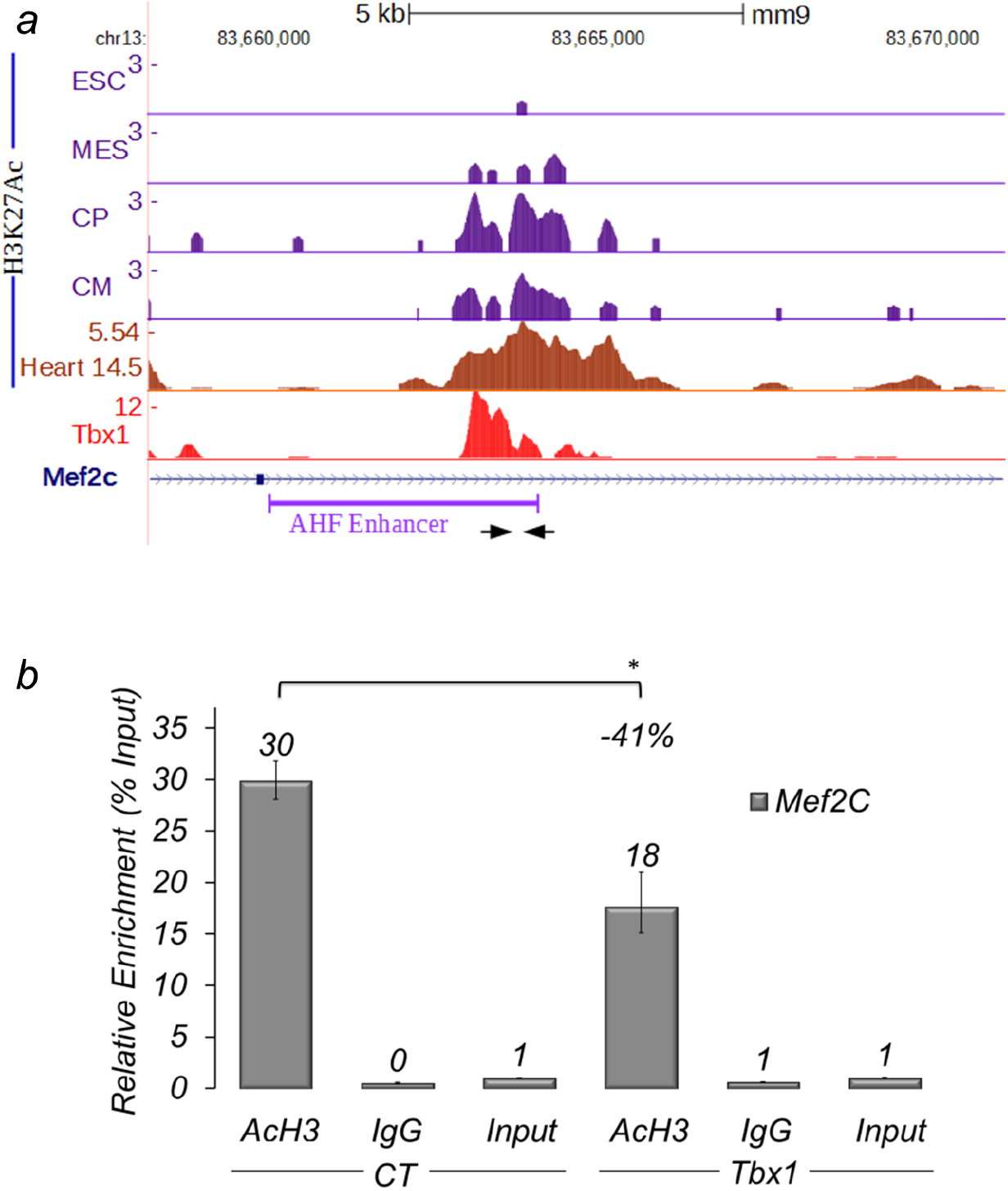
Acetylation of Histone 3 in the AHF enhancer region of Mef2c is reduced by Tbx1 overexpression. (a) Examples of ChIP-seq data profiles as shown using the UCSC genome browser in the genomic region containing the AHF-Enhancer of *Mef2c* (GeneBank AY324098) (a): H3K27Ac profiles at four stages of cardiomyocyte differentiation of mouse embryonic stem cells: undifferentiated embryonic stem cells (ESC), cells expressing mesodermal markers (MES), Cardiac Precursors (CP), cardiomyocytes (CM); H3K27Ac ChIP-seq data profiles in E14.5 heart tissue; Tbx1 ChIP-seq data profiles. Arrows indicate the primers used for real-time PCR amplification. (b) q-ChIP assay of AcH3 on differentiating C2C12 cells (Day 3) transfected with an empty vector (EV) or with a vector over-expressing Tbx1 (Tbx1), followed by quantitative real-time PCR. Values are from 3 experiments (mean± S.D.). Asterisks indicate statistically significant difference (P-value less than 0.05).

These results suggest that Tbx1 contrasts *Mef2c-AHF* enhancer activity by negatively regulating its H3 acetylation.

### Loss of Tbx1 is associated with expansion of transgenic Mef2c-AHF-Cre expression in vivo

To determine whether Tbx1 represses the activity of the Mef2c AHF enhancer in vivo, we used the transgenic Mef2c-AHF-Cre line (Verzi *et al*., 2005), which carries the AHF enhancer that drives the expression of Cre recombinase. The activity of the enhancer was detected using immunofluorescence with an anti-Cre antibody on embryo sections. We tested Mef2c-AHF-Cre;7bx1^+/+^ (control), Mef2c-AHF-Cre;Tbx1^+/-^ (heterozygous), and Mef2c-AHF-Cre;Tbx1^flox/-^ embryos (homozygous null in the Mef2c-AHF-Cre expression domain, and heterozygous elsewhere). We found that at E9.5 (20 somites), Mef2c-AHF-Cre was expressed in the core mesoderm of the 1^st^ and 2^nd^ pharyngeal arches (PAs), in the cardiac outflow tract (OFT), in the SHF (mostly anterior) and in a posterior-lateral cell population of the splanchnic mesoderm at the level of the cardiac inflow tract inlet, in continuity with the SHF (Fig. 3). Cre expression in the more anterior domains (PAs, OFT, and SHF) was comparable between control and mutant embryos (Fig. 3). However, more posteriorly, Cre expression was expanded in both heterozygous and homozygous mutants, albeit more evident in the latter (Fig. 3). The region of expanded Mef2c-AHF-Cre expression is well within the *Tbx1* expression domain at this stage (Supplementary Fig. 3). Thus it is possible that Tbx1 regulates the enhancer directly.

**Figure 3.**
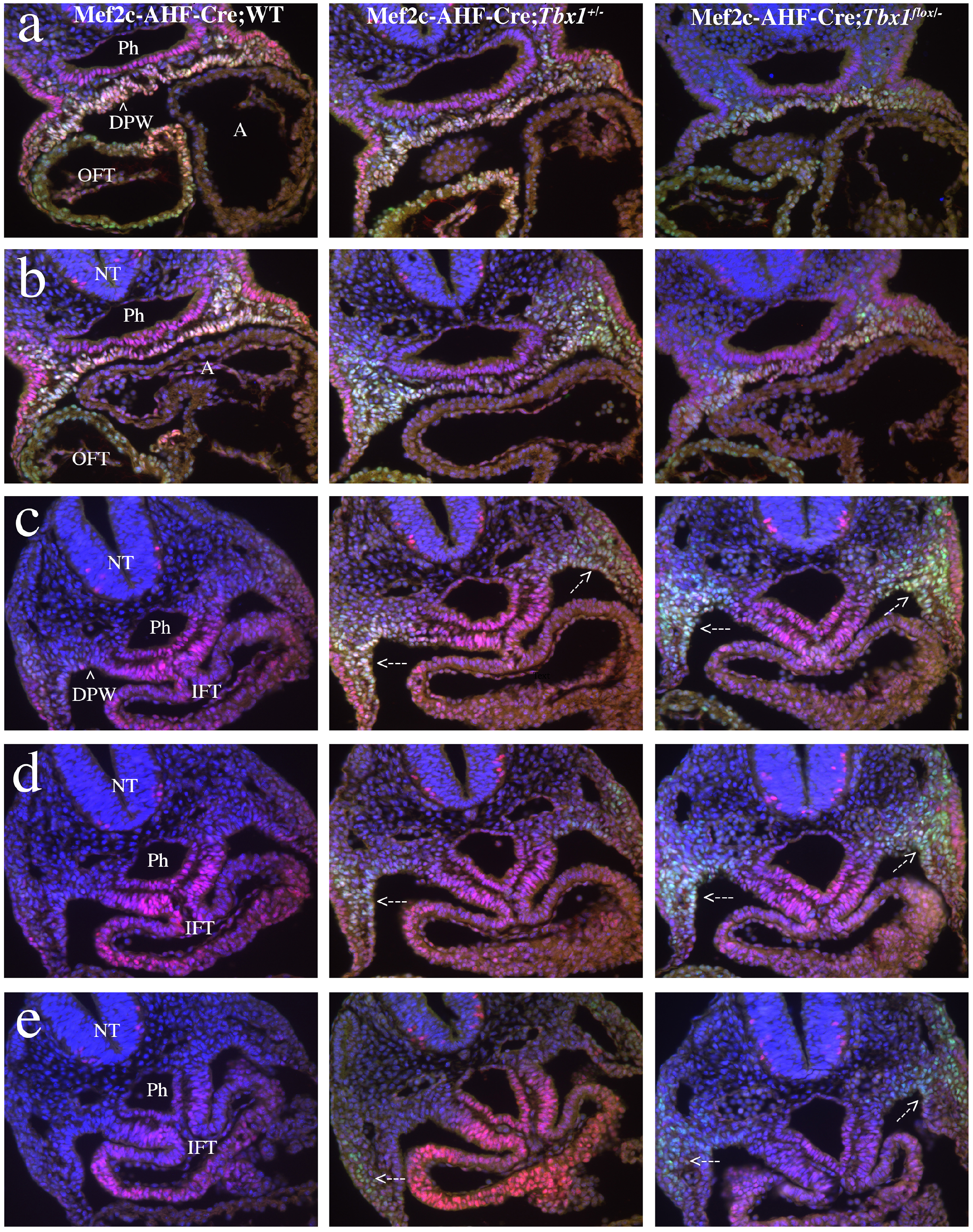
Tbx1 regulates the Mef2c AHF enhancer in vivo. Immunofluorescence of transverse sections of E9.5 (20 somites) embryos with the genotype indicated, using antibodies anti Cre (green) and anti Isl1 (red). The first row (a) of sections correspond to a level immediately below the outflow tract. The second row (b) is just anteriorly to the inflow tract, while the third fourth and fifth rows (c-e) correspond to the inflow tract. Note that the Cre signals in rows a and b are comparable in the three genotypes, while in rows c, d, and e, reduced dosage of *Tbx1* is associated with expansion of Cre expression (arrows). We examined 3 embryos per genotype. Supplementary Figure 3 shows *Tbx1* gene expression in similar sections. A: atrium; DPW: dorsal pericardial wall (arrowhead); IFT: inflow tract; NT: neural tube; OFT: outflow tract; Ph: pharynx;

### Increased expression of MEF2C in 22q11.2DS-derived hiPSCs upon cardiac differentiation

To determine whether the above observations are also relevant for human cells, we have generated induced pluripotent stem cells (hiPSCs) from a 22q11.2DS patient and from an unrelated healthy donor (see cartoon in Fig. 4). hiPSC lines were characterized as shown in Figs. 5 and 6). Two hiPSC clones were differentiated into cardiac progenitors using a previously described protocol (Leschik *et al*., 2008; Moretti *et al*., 2010). We measured *TBX1* expression and found that it peaked at day 4 of differentiation (Fig. 7a), when cells also expressed other markers of cardiac progenitors (Fig. 7b). 22q11.2DS-derived cell lines expressed a lower level of *TBX1*, as the deletion removes one copy of the gene (Fig. 7a).

**Figure 4.**
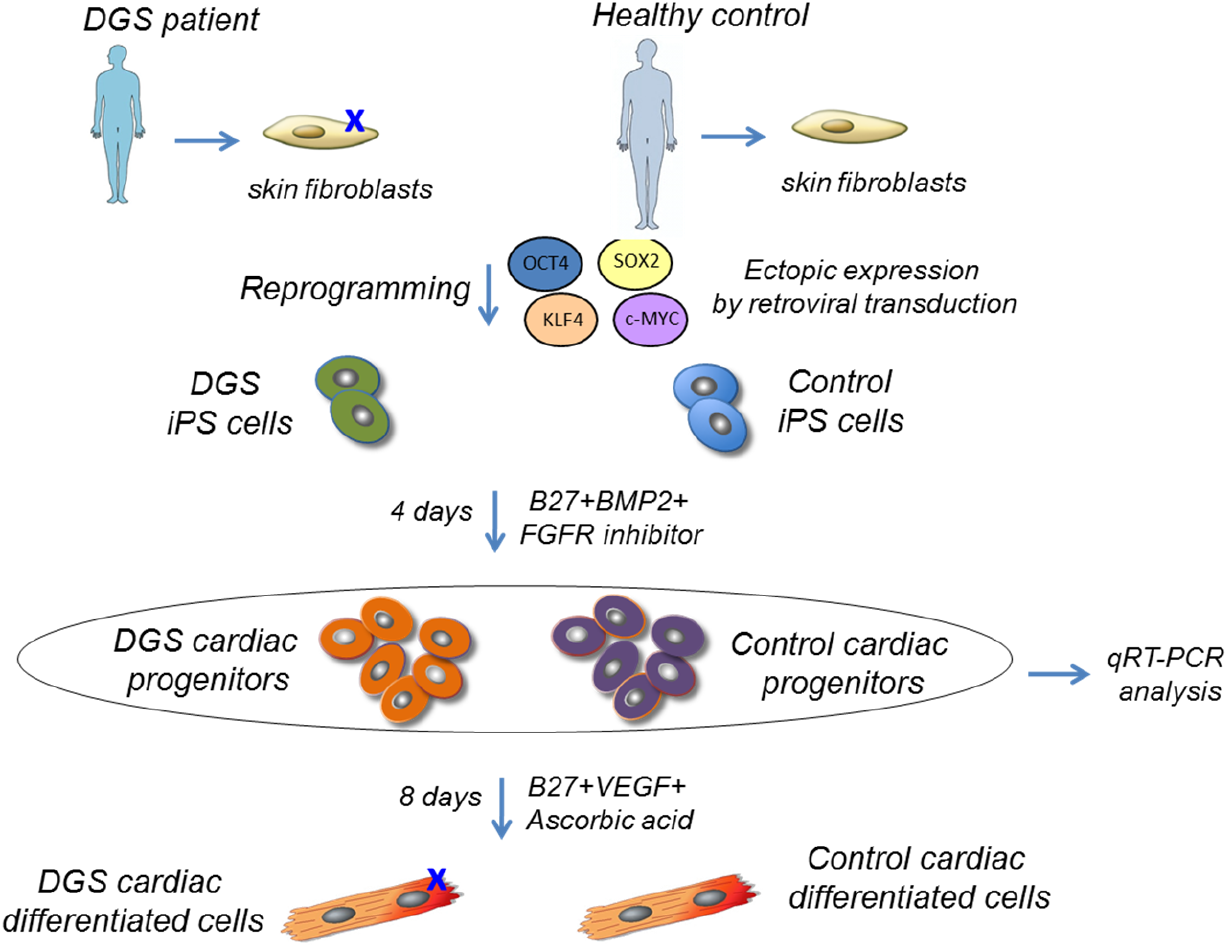
Experimental design for the generation and cardiac differentiation of control and 22q11.2DS induced pluripotent stem cells (iPSCs). Human iPSCs were generated by overexpression of retroviral transgenes for *OCT3/4, SOX2* and *KLF4* in primary skin fibroblasts from a 22q11.2DS patient and from an unrelated healthy donor. Two iPSCs clones from each individual were differentiated in cardiac progenitors and analyzed by quantitative RT-PCR (qRT-PCR).

**Figure 5.**
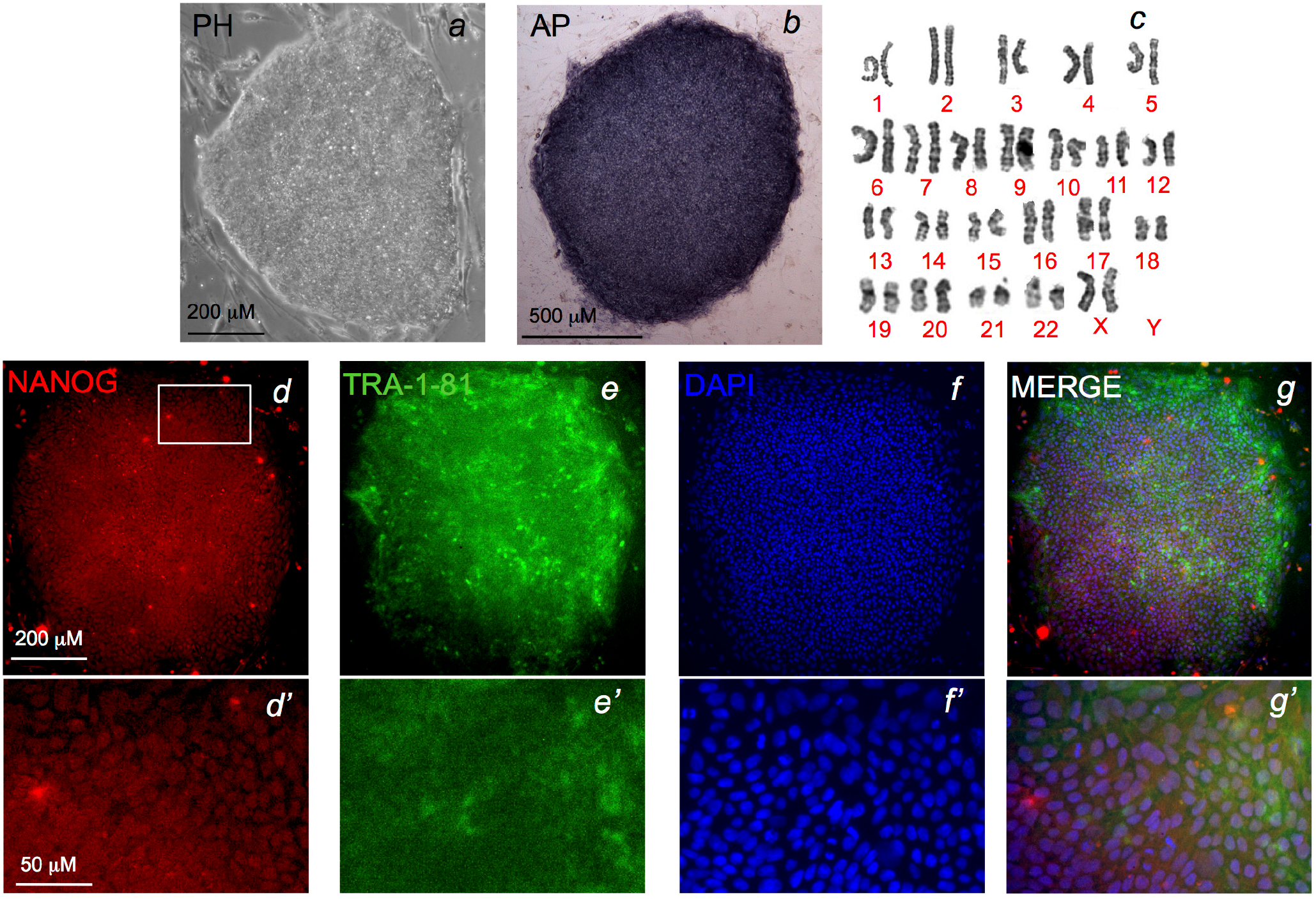
Generation and characterization o 22q11.2DS patient-derived iPSCs. Images of colonies from a representative 22q11.2DS iPSC clone in bright field (a) and after staining for alkaline phosphatase (AP) activity (b). Scale bar, 500 and 200 *μ*m respectively. c, Karyotyping of a representative 22q11.2DS iPSC clone. d-f’, Immunofluorescence analysis of pluripotency markers NANOG (d,d’) and TRA1-81 (e, e’) in a representative iPSC clone. DNA is stained with DAPI (f,f’). Scale bar, 200 *μ*m. d’, e’ and f’ are magnification of d, e and f respectively. Scale bar, 50 *μ*m.

**Figure 6.** Assessment of pluripotency in control and 22q11.2DS iPSCs. a, Quantitative RT-PCR (qRT-PCR) for expression of retroviral transgenes in two iPSC clones from a control individual (CTRcl1 and CTRcl2) and two from a 22q11.2DS patient (22q11.2DScl1 and 22q11.2DScl2). Expression values are relative to the corresponding primary skin fibroblasts (PSF) infected with the four retroviruses *OCT3/4, SOX2* and *KLF4* (Infected prmary skin fibroblasts, IPSF), normalized to *GAPDH*, and presented as mean ± s.e.m., n=3. b, qRT-PCR analysis of endogenous genes associated with pluripotency (*c-MYC, KLF4, OCT3/4, SOX2, NANOG, REX1*, and *TDGF1*) in the two control (CTRcl1 and CTRcl2)and two 22q11.2DS iPSC clones (22q11.2DScl1 and 22q11.2DScl2). Expression values are relative to corresponding PSF, normalized to GAPDH, and presented as mean ± s.e.m., n=3. c, qRT-PCR analysis of markers of the three germ layers, endoderm (*PDX1, SOX7*, and *AFP*), mesoderm (*CD31, DES, ACTA2, SCL, MYL2*, and *CDH5*), and ectoderm (*KRT14, NCAM1, TH*, and *GABRR2*) in embryoid bodies (EBs) at day 21 of differentiation from the two control (CTRcl1 and CTRcl2 EBs) and two 22q11.2DS iPSC clones (22q11.2DScl1 and 22q11.2DScl2 EBs). Expression values are relative to the corresponding undifferentiated iPSC clones, normalized to *GAPDH*, and presented as mean ± s.e.m., n=3.

**Figure 7.**
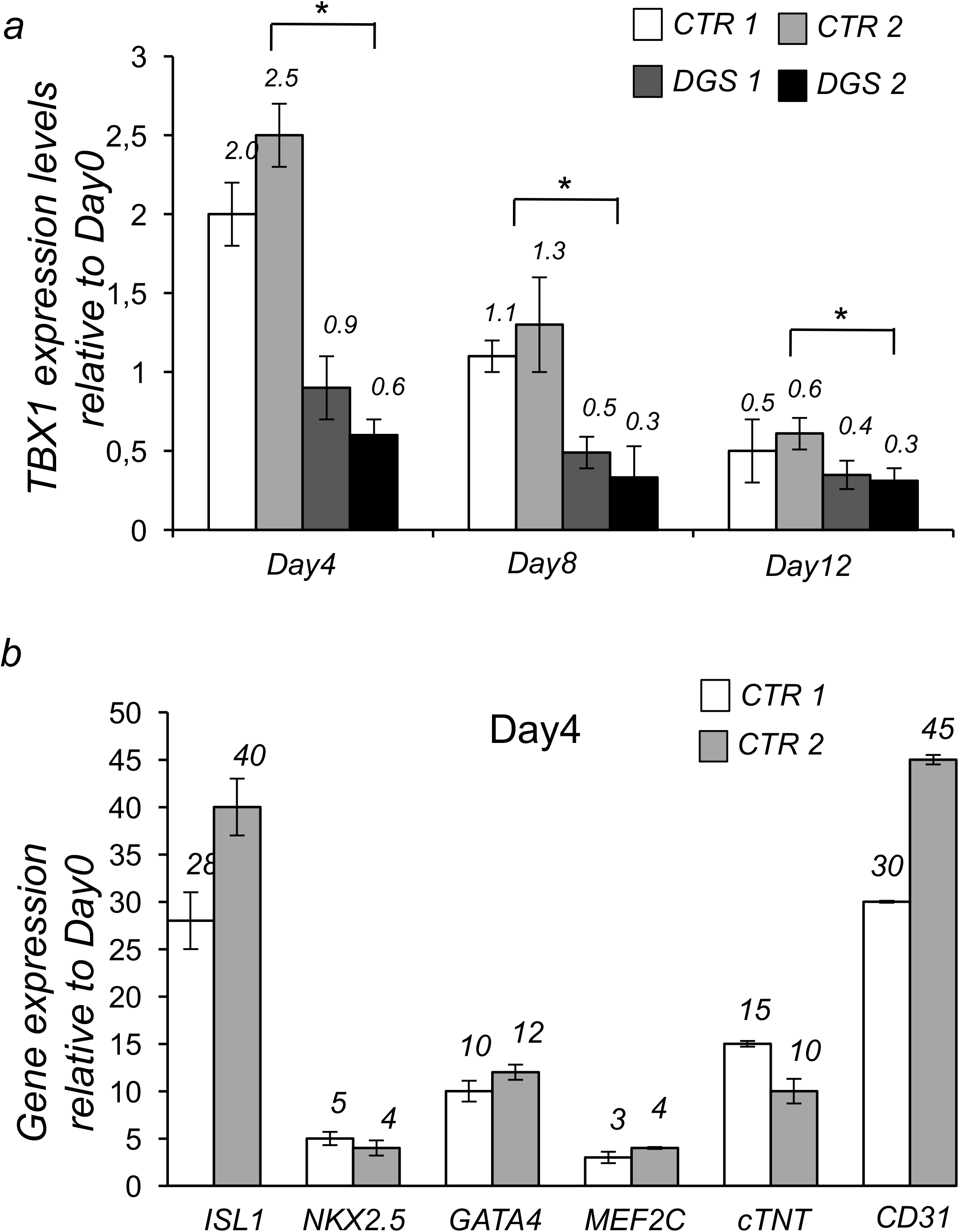
Analysis of TBX1 expression in control and 22q11.2DS iPSC-derived cardiac cells. a, Quantitative RT-PCR (qRT-PCR)-based TBX1 expression profile in two iPSC clones from a control individual and two from a 22q11.2DS patient during cardiac differentiation. b, qRT-PCR-based comparison of expression of the genes indicated in the two control iPSC clones after 4 days BMP2/FGFR-inhibitor treatment (Day4). n=3,*P<0.05 Vs. Day 0.

We measured the expression of *MEF2C* and *GATA4*, which is another target of Tbx1 (Liao *et al*., 2008; Pane *et al*., 2012), by qRT-PCR at days 4, 8, and 12 of differentiation and found that both genes are up regulated in patient-derived cells at all differentiation points tested, compared to control cells (Fig. 8a-b).

**Figure 8.**
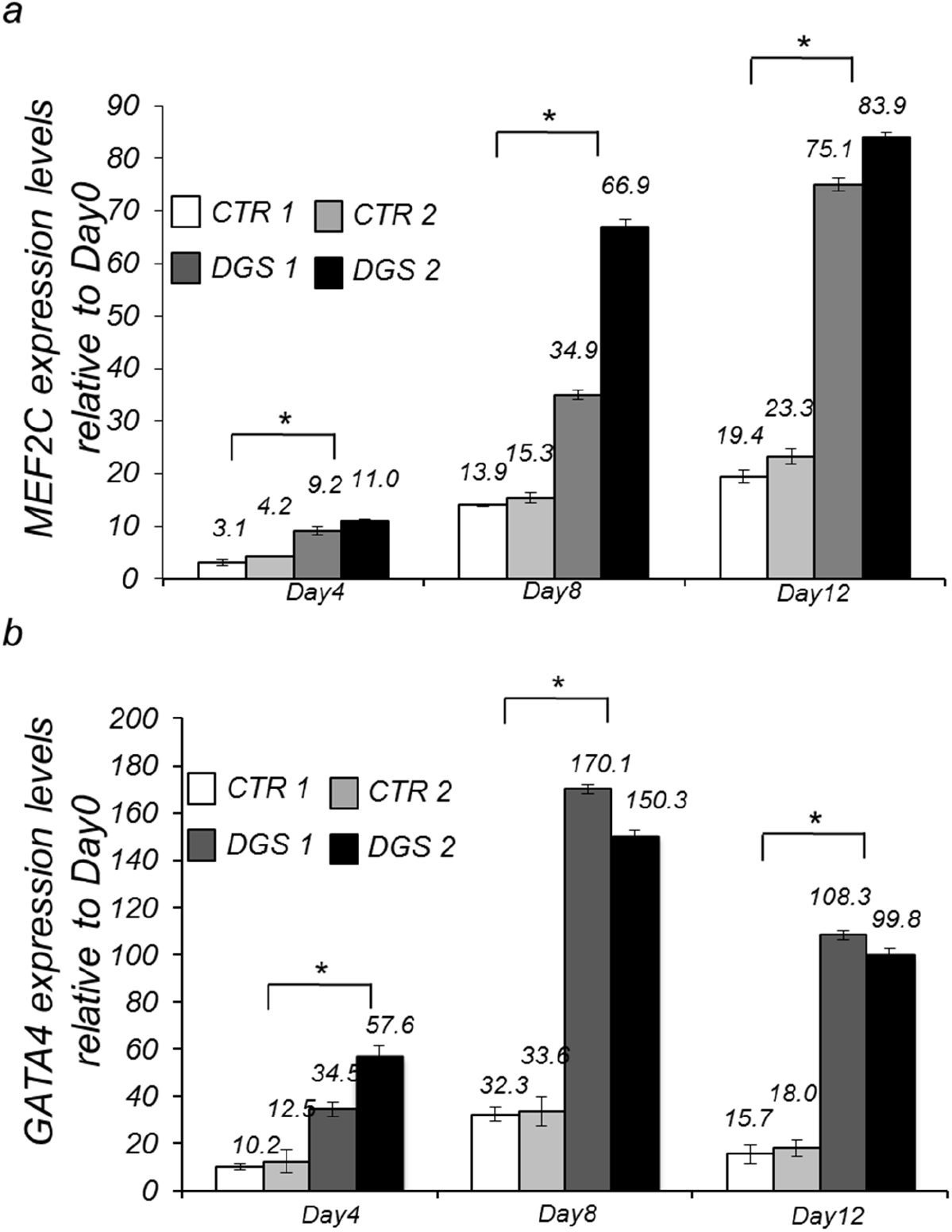
Analysis of MEF2C and GATA4 expression in control and 22q11.2DS iPSC-derived cardiac cells. Quantitative RT-PCR (qRT-PCR)-based MEF2C (a) and GATA4 (b) expression in two iPSC clones from a control individual and two from a 22q11.2DS patient at Day4, Day8 and Day12 of cardiac differentiation. n=3,*P<0.05 Vs. Day 0.

Next, we performed q-ChIP assays on the AHF enhancer homologous region of the *MEF2C* gene and on the *GATA4* promoter. Results demonstrated that H3Ac enrichment is significantly higher in 22q11.2DS cells compared to controls for both genes (Fig. 9).

**Figure 9.**
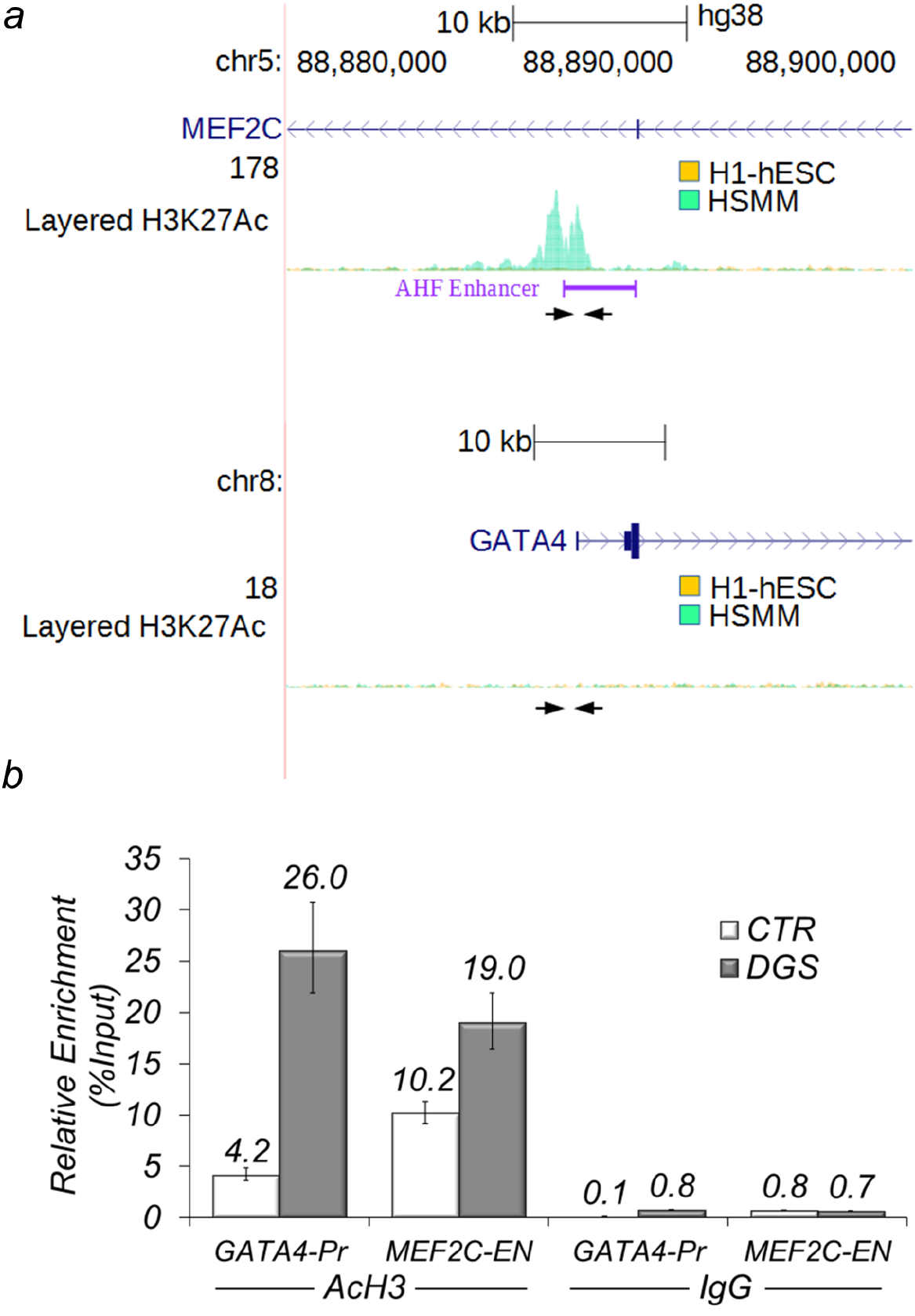
Acetylation of Histone 3 in MEF2C and GATA4 loci is enhanced in differentiating 22q11.2DS iPS cells. (a,b) Localization of primer pairs (arrows) used for ChIP experiments with human cells on *MEF2C* and *GATA4* genes. ″AHF enhancer″ indicates a region of high similarity with the murine AHF enhancer. For reference, we report ENCODE ChIP-seq data for H3K27Ac for two cell types, undifferentiated human embryonic stem cells (H1-hESC) and human skeletal muscle myoblasts (HSMM). The GATA4 promoter region is not acetylated in these two cell types. (c) ChIP assays with an antibody anti-AcH3 in differentiating control and 22q11.2DSiPS cells followed by quantitative real-time PCR.

### Overall, our data indicate that Tbx1 functions as a transcriptional repressor of the *Mef2c*

AHF enhancer in mouse and human cells and in vivo, in mouse embryos. In embryos, we noted that the response of the enhancer is *Tbx1* dosage-dependent and is restricted to the splanchnic mesoderm, in a posterior-lateral domain, indicating context-dependent regulation. The AHF enhancer integrates the actions of several critical transcription factors (Dodou *et al*., 2004), e.g. Isl1, Nkx2-5, Gata4, and also Tbx1. Interpreting the combinatorial code governing its regulation is of interest for the understanding of SHF development. We have previously shown that in cultured cells, Tbx1 inhibits Gata4-mediated activation of the AHF enhancer, but not Isl1-mediated activation (Pane et al., 2012). Interestingly, here we noted that the Mef2c-AHF-Cre expanded expression domain in mutant embryos is not stained with our Isl1 immunofluorescence, while most of the other Mef2c-AHF-Cre expression domains are Isl 1 - positive. In the future, it would be of interest to establish whether Isl 1 represents a ″protective factor″ against Tbx1-dependent suppression.

Ripply3 binds Tbx1 and inhibits its transcriptional activity in cell culture experiments (Okubo *et al*., 2011). In addition, and consistently, Ripply3 turns Tbx1 into a transcriptional repressor in Xenopus (Janesick *et al*., 2012). Ripply proteins interact with Groucho/TLE proteins which, in turn, can repress gene expression using various mechanisms, for example induction of chromatin compaction (Sekiya and Zaret, 2007) or direct interaction with HDACs, including HDAC1 (Brantjes *et al*., 2001; Jennings and Ish-Horowicz, 2008). Groucho also interacts with Tbx15, Tbx18, and Tbx20 (Farin *et al*., 2007; Kaltenbrun *et al*., 2013), but Tbx1 lacks the Groucho interaction domain (Farin *et al*., 2007). In addition, Ripply3 expression is restricted to the mouse pharyngeal endoderm (Okubo *et al*., 2011), so it is unlikely to play a role on the observed expansion of the Mef2c-AHF-cre domain in the splanchnic mesoderm. Our data showed that Tbx1 does not interact directly with HDAC1, suggesting the existence of molecular intermediates responsible for driving histone deacetylation. Of particular interest is the location in which Tbx1 effects this function as it may be related to sorting anterior vs. posterior heart field, and/or affect cardiomyocyte progenitors contribution to the OFT. Future studies should be directed to the identification of molecular intermediate(s) operating in the splanchnic mesoderm and affecting the transcriptional activity of Tbx1 in this region.

## Acknowledgments

We thank Dr. Thomas Zwaka and his lab members for invaluable help in establishing methods for hiPSC generation and characterization. We acknowledge the support of the Integrated Microscopy Facility of the Institute of Genetics and Biophysics and the Cell Culture Facility of TIGEM, Pozzuoli, Italy. This work was made possible by funding from the Italian Telethon Foundation (GGP14211), from the Fondation Leducq (TNE 15CVD01), and from the MIUR/FAREBIO grant to AB.

## Supplementary Table 1

Sequence of oligonucleotides used in this study

## Supplementary Figure 1

Histogram showing the results of Q-ChIP analyses using anti H3K27Ac antibodies on C2C12 cells treated with non-targeting siRNA (*Tbx1*^WT^) or with Tbx1-targeted siRNA (*Tbx1^KD^*) on the same locus as the one tested in Fig. 2. Enrichment is shown as percentage of input. Results are the mean of two biological replicates (error bars indicate s.e.m.).

## Supplementary Figure 2

Western blot analysis of representative immunoprecipitation experiment using protein extracts from Tbx1:3xHA-transfected C2C12 cells. Immunoprecipitation of nuclear extracts was performed using anti-HA antibodies or IgG (control). Top panel: control WB using antiHA antibodies. Lower panel: WB with anti-HDAC1 antibodies. Immunoreactivity is found in the FT and input samples but not in the immunoprecipitated samples (E11 and El2).

The experiment has been repeated 5 times. FT: flow-through; E11: Elution 1; El2: Elution 2; CE: cytoplasm extract; NE: nuclear extract.

## Supplementary Figure 3

Transverse sections of a *Tbx1*^*/lacZ^ embryo at E9.5 (22 somites) after staining with the β-gal chromogenic substrate salmon-gal. Sections a-e are at similar levels as sections a-e shown in Fig. 3. Bar indicates 100*μ*m. f-g: left and right sides of the whole-mount stained embryo from which sections a-e were derived. h: WT littermate with negative staining.

DPW: dorsal pericardial wall; IFT: inflow tract; Ph: pharynx.

